# CHD4 is recruited by GATA4 and NKX2-5 silencers to repress non-cardiac gene programs in the developing heart

**DOI:** 10.1101/2021.10.28.466285

**Authors:** Zachary L. Robbe, Wei Shi, Lauren K. Wasson, Caralynn M. Wilczewski, Xinlei Sheng, Austin J. Hepperla, Brynn N. Akerberg, William T. Pu, Ileana M. Cristea, Ian J. Davis, Frank L. Conlon

## Abstract

The Nucleosome Remodeling and Deacetylase (NuRD) complex is one of the central chromatin remolding complexes that mediate gene repression. NuRD is essential for numerous developmental events, including heart development. Clinical and genetic studies have provided direct evidence for the role of chromodomain helicase DNA-binding protein 4 (CHD4), the catalytic component of NuRD, in congenital heart disease (CHD), including atrial and ventricular septal defects. Further, it has been demonstrated that CHD4 is essential for mammalian cardiomyocyte formation and function. A key unresolved question is how CHD4/NuRD is localized to specific cardiac targets genes, as neither CHD4 nor NuRD can directly bind DNA. Here, we coupled a bioinformatics-based approach with mass spectrometry analyses to demonstrate that CHD4 interacts with the core cardiac transcription factors GATA4, NKX2-5 and TBX5 during embryonic heart development. Using transcriptomics and genome-wide occupancy data, we have characterized the genomic landscape of GATA4, NKX2-5 and TBX5 repression and defined the direct cardiac gene targets of GATA4-CHD4, NKX2-5-CHD4 and TBX5-CHD4 complexes. These data were used to identify putative cis-regulatory elements regulated controlled by these complexes. We genetically interrogated two of these silencers in vivo, *Acta1* and *Myh11*. We show that deletion of these silencers leads to inappropriate skeletal and smooth muscle gene mis-expression, respectively, in the embryonic heart. These results delineate how CHD4/NuRD is localized to specific cardiac loci and explicates how mutations in the broadly expressed CHD4 protein lead to cardiac specific disease states.

## Introduction

Transcriptional repression is mediated by broadly expressed multiprotein complexes that modify and remodel chromatin. Repression by these complexes is achieved by alteration of chromatin states through direct DNA-sequence-specific binding of a given complex or by recruitment of the complex to defined loci via interactions with tissue-specific transcription factors. The Nucleosome Remodeling and Deacetylase (NuRD) complex represses gene expression and is essential from fly to human. In most cases, the combined histone deacetylase and ATP-dependent chromatin remodeling helicase of NuRD combine to modulate chromatin states at target genes (Wade et al. 1998; Xue et al. 1998; Zhang et al. 1998; Wade et al. 1999). Components of the NuRD complex differ but invariably contain either of the ATP-dependent chromodomain helicases CHD3 or CHD4 (Watson et al. 2012; Joshi et al. 2013; Low et al. 2016; Zhang et al. 2016; Hoffmeister et al. 2017; Farnung et al. 2020). The crucial nature of CHD4 has been established through clinical studies that have shown mutations in CHD4 lead to congenital heart disease (Zaidi et al. 2013; Homsy et al. 2015; Sifrim et al. 2016; Waldron et al. 2016; Weiss et al. 2016). In addition, genetic studies in mouse have demonstrated that cardiac conditional Chd4 null mutants die at mid-gestation and that loss of CHD4-mediated repression leads to misexpression of fast skeletal and smooth muscle myofibril isoforms, cardiac sarcomere malformation and early embryonic lethality (Gomez-Del Arco et al. 2016; Wilczewski et al. 2018).

Though CHD4 is essential for heart development, and its disease relevance has been shown, one of the central unanswered questions regarding the complex’s regulatory mechanism is how CHD4/NuRD is recruited to specific cardiac gene loci, given that neither CHD4 nor NuRD bind DNA. Using a genomic approach, we identified overrepresented DNA motifs recognized by several core cardiac transcription factors at CHD4-bound genomic regions. Using parallel reaction monitoring mass spectrometry (PRM nLC-MS/MS) (Picotti et al. 2013; Federspiel et al. 2019), we confirmed that CHD4 interacts *in vivo* in mid-gestation hearts with GATA4, NKX2-5 and TBX5. Further analysis revealed that CHD4 and GATA4, NKX2-5 and TBX5 converge on regulatory elements genome-wide associated with transcriptional repression. These findings imply an important dual regulatory role for these critical cardiac transcription factors.

We have used our data to map putative cardiac cis-regulatory elements (CREs) regulated through GATA4-CHD4, NKX2-5-CHD4, and TBX5-CHD4 complexes. Deletion of a putative CREs at the skeletal muscle *Acta1* gene that contains an NKX2-5 binding site led to inappropriate *Acta1* expression in the fetal heart. Similarly, deletion of a CHD4 CRE of the smooth muscle *Myh11* gene containing a GATA4 binding site led to *Myh11* mis-expression in the fetal heart. Collectively, our results demonstrate that CHD4/NuRD is recruited to defined loci through a core set of cardiac transcription factors to repress inappropriate gene expression in the developing heart.

## Results

### CHD4 interacts with the core cardiac transcription factors GATA4, NKX2-5, and TBX5

One of the central issues regarding the mechanism and function of the CHD4/NuRD complex involves the recruitment of CHD4/NuRD to specific loci, given that neither CHD4 nor NuRD bind DNA directly (Figure 1A). To address these issues, we performed motif discovery of CHD4 ChIP-seq data sets obtained from E10.5 hearts (Wilczewski et al. 2018) (GEO:GSEI090I2). Our analyses revealed a striking abundance of significantly over-represented cardiac transcription factor consensus motifs in CHD4-bound regions (Figure 1B). These sequences included binding sites for the core cardiac transcription factors GATA4, NKX2-5, and TBX5, each of which is required for cardiac development and, when mutated, causes human heart disease (Durocher et al. 1997; McCulley and Black 2012; Baban et al. 2014; Akerberg et al. 2019; Jumppanen et al. 2019). Moreover, mice homozygous null for *Tbx5, Gata4* and *NKX2-5* display heart defects at E10.5, the same developmental stage that requires *Chd4* (Bruneau et al. 2001; Stennard et al. 2003; Watt et al. 2004; Mori et al. 2006; Maitra et al. 2009; Terada et al. 2011; Zhou et al. 2015; Gomez-Del Arco et al. 2016; Wilczewski et al. 2018).

**Figure 1.**
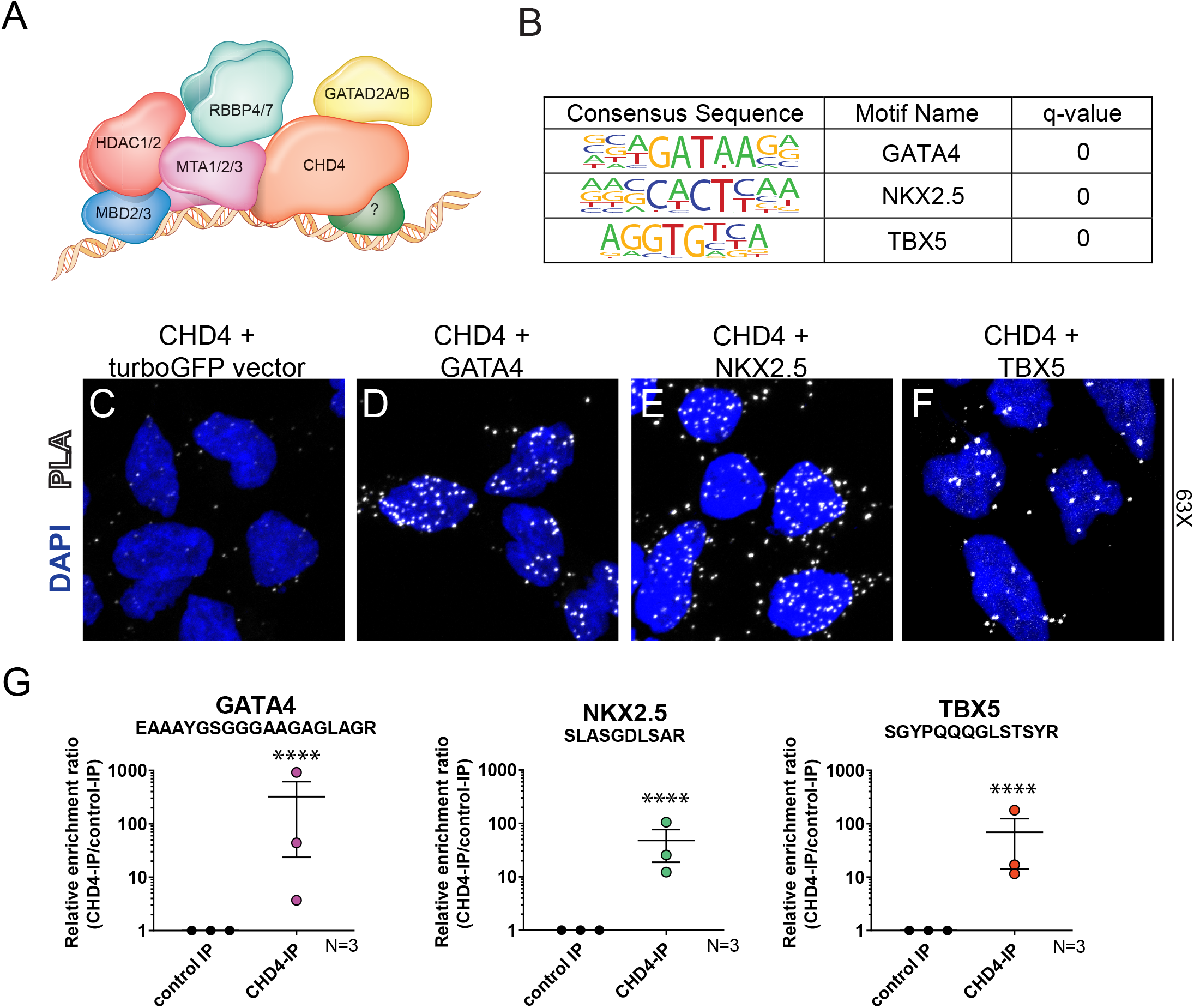
CHD4 Interacts with Core Cardiac TFs: GATA4, NKX2-5, and TBX5. (A) Schematic of the Nucleosome Remodeling and Deacetylase (NuRD) complex localizing to target loci through interaction with a tissue-specific co-factor. (B) Predicted co-occupancy of CHD4 and GATA4, NKX2-5, and TBX5 through significant overrepresentation of known binding motifs of each factor in CHD4 bound genomic regions. q values = 0.0000 generated from FDR-corrected P value based on the cumulative hypergeometric distribution. (C-F) CHD4 associates with GATA4, NKX2-5, and TBX5 through proximity ligation assay. Each image shown as a max intensity z-stack projection using a 63xoil magnification objective. Presence of PLA signal (white dots) denotes close physical proximity of target proteins and represents a physical interaction. CHD4 and turboGFP alone (C) result in a lower frequency and intensity of PLA signal than CHD4-GATA4 (D), CHD4-NKX2-5 (E), or CHD4-TBX5 (F). (G) CHD4 complexes affinity purified from wild-type E10.5 mouse embryonic hearts show a significant enrichment for peptides belonging to GATA4, TBX5, or NKX2-5 when compared to purified GFP complexes, as determined by PRM-MS quantification and Student’s t test. Data shown as mean ± SEM of triplicates, n = 3, with each replicate of >28 pooled embryonic hearts per IP. **** P value < 0.0001.

We confirmed CHD4 co-localization with GATA4, NKX2-5 and TBX5 by proximity ligation assays (PLA). CHD4 was expressed with either GATA4, NKX2-5, TBX5, or as a negative control, turboGFP (Figures 1C-F). In support of our genomic analyses, we find that CHD4 interacts with GATA4, NKX2-5, and TBX5, and in all three cases, the interaction occurs predominantly in the nucleus.

To elucidate if CHD4 interacts with either GATA4, NKX2-5 or TBX5 in heart tissue *in vivo*, we performed parallel reaction monitoring MS (PRM LC-MS/MS) on immuno-affinity purified cardiac E10.5 CHD4 interactomes (Figure 1G). Consistent with previous findings (Waldron et al. 2016), PRM LC-MS/MS analysis of E10.5 CHD4 endogenous interactomes revealed CHD4 in complex with TBX5. These analyses also identified that CHD4 is in complex with GATA4 and NKX2-5 in E10.5 hearts (Figure 1G). To our knowledge, this is the first report of an endogenous interaction between CHD4 and NKX2-5 during heart development. Together, our data establish that CHD4 complexes *in vivo* with GATA4, NKX2-5 and TBX5 during a time point when CHD4 is essential for vertebrate heart development.

### GATA4, NKX2-5 and TBX5 are co-bound with CHD4

To determine if GATA4, NKX2-5, and TBX5 binding sites were prevalent in CHD4 ChIP-seq peaks, we overlapped GATA4, NKX2-5, and TBX5 ChIP-seq datasets (Akerberg et al. 2019) with CHD4 ChIP-seq peaks. We observe that the majority of CHD4-bound peaks are also bound by GATA4, NKX2-5, and/or TBX5 (Figure 2A).

**Figure 2.**
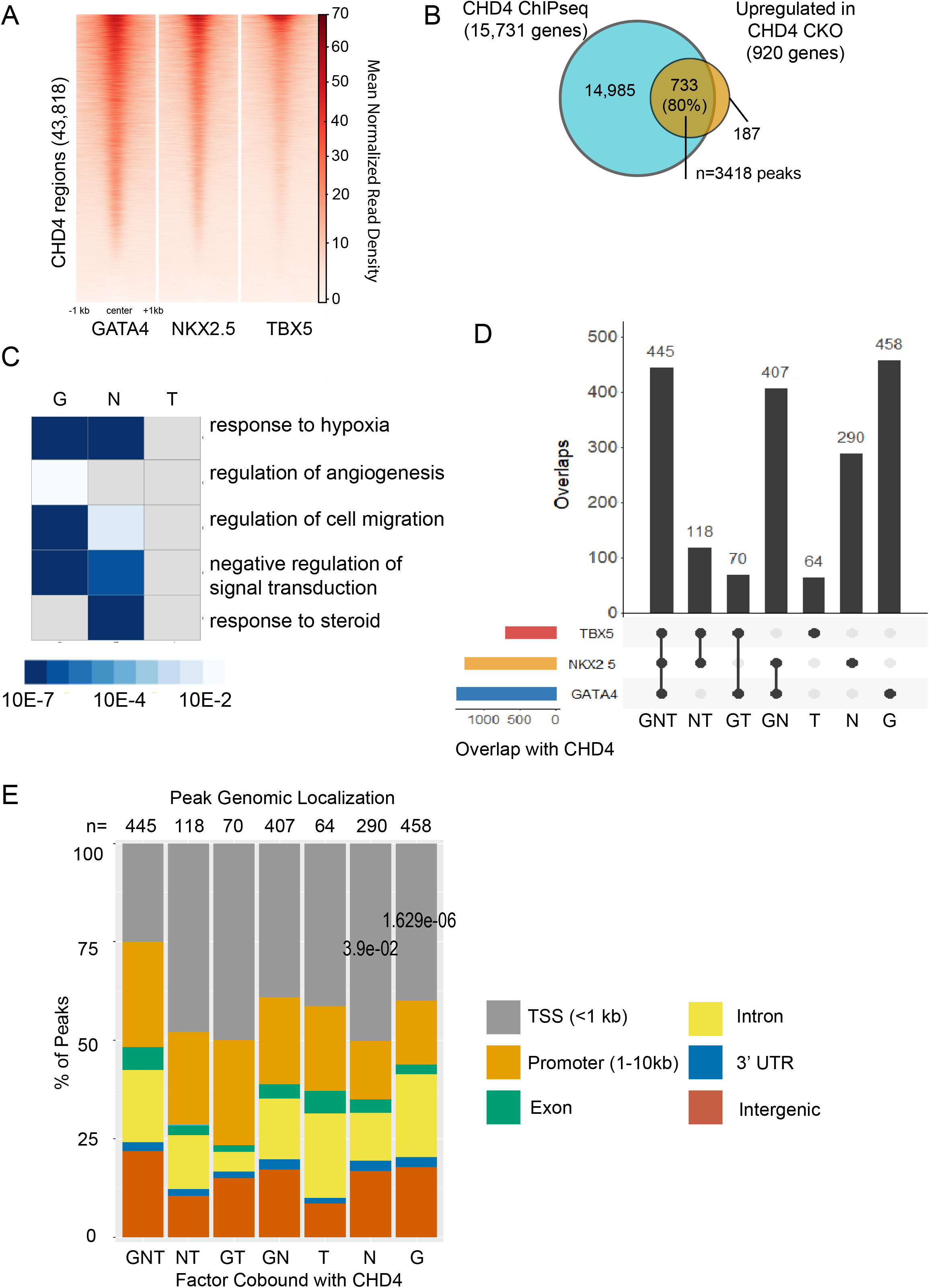
CHD4 Co-occupies Directly Repressed Loci with GATA4, NKX2-5, and TBX5 (A) Heatmap visualizing density of ChIP-seq signal of each transcription factor (GATA4, NKX2-5, and TBX5) across genomic regions occupied by CHD4 in the developing heart. (B) Overlap of genes bound by CHD4 and upregulated in Chd4-null hearts. 733/920 (80%) of genes upregulated in hearts devoid of CHD4 are also bound by CHD4 at E10.5. These 733 genes are termed CHD4-repressed genes and are associated with 3418 peaks. (C) Biological processes overrepresented in regions co-occupied and repressed by CHD4-GATA4, CHD4-NKX2-5, and CHD4-TBX5. The heatmap is presented as the -log Bonferroni corrected P value. Gray boxes represent Biological Processes with a non-significant P-value for that group. (D) Upset plot displaying intersection of ChIP-seq peaks shared between CHD4 and GATA4, NKX2-5 and TBX5, respectively. Each column represents a different possible overlap of the data, with the totals defined with the horizontal-colored bars. CHD4 peaks that did not overlap with any transcription factor are not depicted in this plot (E) Stacked bar plot visualizing the distribution of peaks to their associated gene feature annotated using ChIP-Seeker. Columns represent transcription factor peaks associated with CHD4-repressed genes or other regions. Statistical significance was calculated using a t-test, comparing each column to GNT.

As we had previously shown that CHD4 interacts with TBX5 to repress non-cardiac gene programs (Waldron et al. 2016), we hypothesized that the interactions with GATA4 and NKX2-5 have a similar consequence. Therefore, we focused on sites within genes that were also bound by CHD4 and that demonstrated increased RNA levels following CHD4 knock out, supportive of a CHD4 repressive activity (Figure 2B) (n=3418 peaks). Gene Ontology analysis of peaks cobound by CHD4 and GATA4, NKX2-5, or TBX5 revealed potential biological differences between the CHD4 interaction with each of these transcription factors. Specifically, CHD4/GATA4 and CHD4/NKX2-5 co-bound peaks were linked to genes associated with hypoxia, angiogenesis, and cell migration. In contrast, CHD4/TBX5-co-bound peaks did not demonstrate this association (Figure 2C).

As there are well-characterized interactions between TBX5, GATA4, and NKX2-5 (Durocher et al. 1997; Jumppanen et al. 2019) we hypothesized that there are genomic loci at which CHD4 binds more than one transcription factor. Of the CHD4-bound peaks in repressed genes, more than half (n=1852) were co-bound by TBX5, GATA4, and/or NKX2-5 (Figure 2D). Furthermore, we found that 445 peaks at CHD4 repressed genes were bound by all three transcription factors in addition to CHD4 (Figure 2D). GATA4 was found most often to be associated with CHD4-repressed regions and was also the most likely to co-bind with CHD4 individually (Figure 2D).

We next determined the genomic localization of the CHD4/transcription factor binding to determine a potential effect on gene expression. We annotated binding peaks with the nearest target gene feature (Figure 2E). These analyses show that GATA4-CHD4, NKX2-5-CHD4 and TBX5-CHD4 are significantly enriched at intergenic, intronic and promoter regions, suggesting that they regulate gene expression, as these sites are associated with CHD4-mediated transcriptional repression(Wilczewski et al. 2018). We next sought to determine if the number of bound transcription factors or their composition affected the magnitude of CHD4 gene repression. To this end, we examined changes in expression in CHD4/GATA-bound, CHD4/NKX2-5-bound, and/or CHD4/TBX5-bound genes (Figure S1). We conclude that the magnitude of CHD4 repression is not dependent upon the identity of the recruiting factor or the number of occupying factors. Thus, CHD4 interaction with GATA4, NKX2-5 and TBX5 is critical for the localization of the complex to target loci, but the composition of the interaction does not affect the extent of transcriptional repression.

### GATA4 is the predominant CHD4 co-factor

Our data infers that, of the three cardiac core transcription factors, GATA4 is the transcription factor most frequently associated with CHD4 (Figure 2D). To determine further the potential role for GATA4 in repression, we mapped GATA4, NKX2-5 and TBX5 ChIP-seq read density across CHD4 repressed regions grouped by number of bound transcription factors (Figure 2A). From the data, we discovered that both NKX2-5 and TBX5 have a lower read density when bound alone than when bound with at least one other factor, the opposite of what is observed for GATA4. We conclude that GATA4 is the predominant cardiac co-factor for recruiting CHD4.

### NKX2-5 recruits CHD4 to repress expression of skeletal actin in the embryonic heart

Strikingly, we found that GATA4 was localized to 82% (1166/1422) of all CHD4-repressed loci. We, therefore, addressed if binding of either NKX2-5 or TBX5 at a single site is sufficient to recruit CHD4 at CHD4 repressed genes. We identified putative silenced enhancers in cardiac tissue focusing on target genes that: 1) contained a single binding site for either NKX2-5 or TBX5, 2) demonstrated binding site conservation between mouse and human, 3) were associated with a single CHD4 peak and, 4) were marked by H3K27me3 (Figure 3)(Consortium 2012). Among genes meeting these four criteria was *Acta1*, the gene that encodes a skeletal actin isoform and is an established CHD4/NuRD target (Wilczewski et al. 2018). In the normal developing heart, cardiac actin *Actc1* is the predominant actin isoform (Mayer et al. 1984) whereas *Acta1*, the skeletal isoform, is expressed at low to undetectable levels. The *Acta1* locus contained a single putative NKX2-5 DNA-binding motif located in the *Acta1* 3’UTR (Figures 4A-C). Thus, the 3’UTR of *Acta1* was postulated to contain a silent enhancer bound by NKX2-5 and CHD4 in E10.5 hearts.

**Figure 3.**
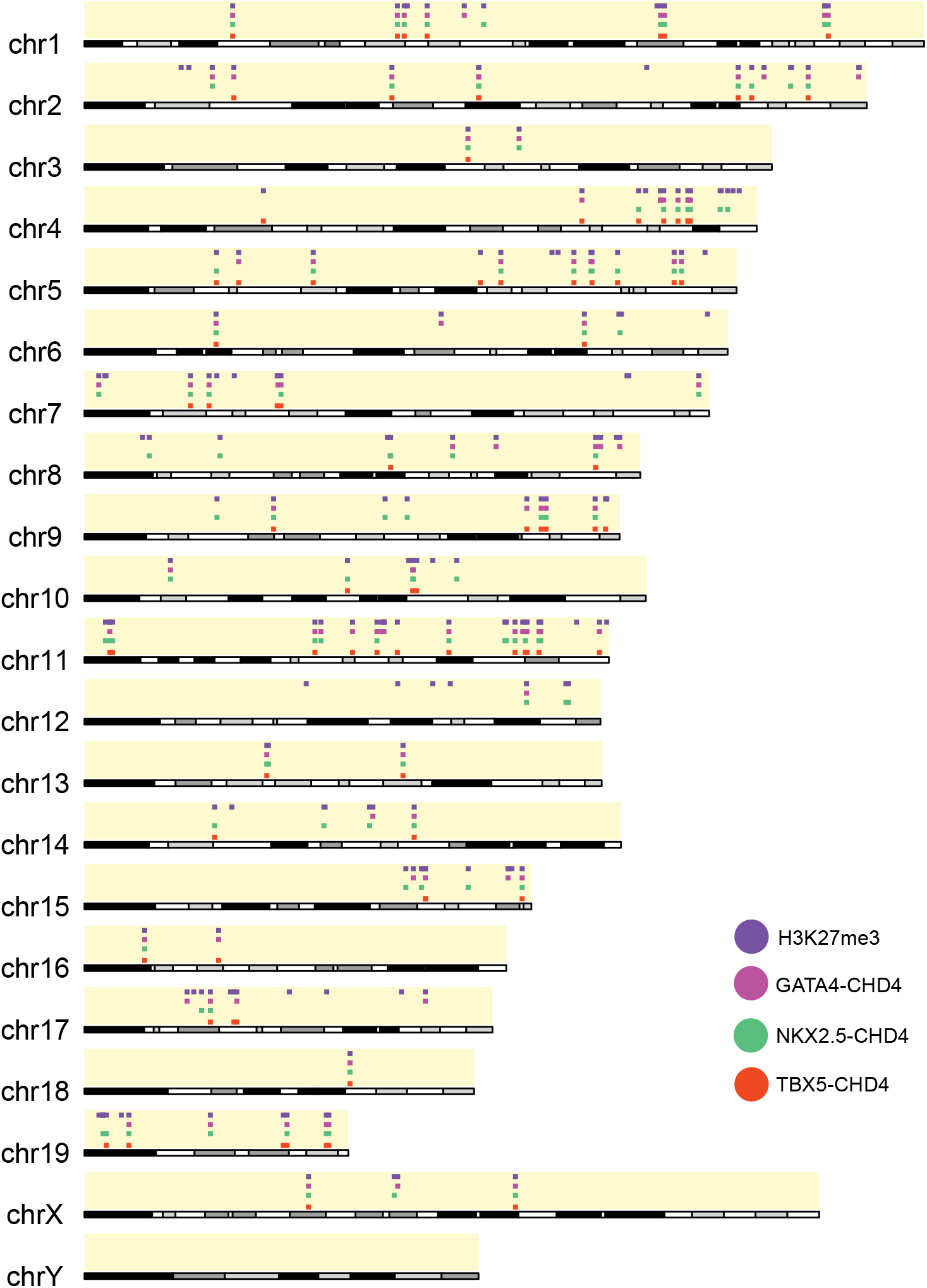
GATA4, NKX2-5, and TBX5 are Associated with H3K27me3 at CHD4-repressed Genes Sites of CHD4 and transcription factor association at CHD4-repressed genes marked with H3K27me3 histone modification at a chromosome level.

**Figure 4.**
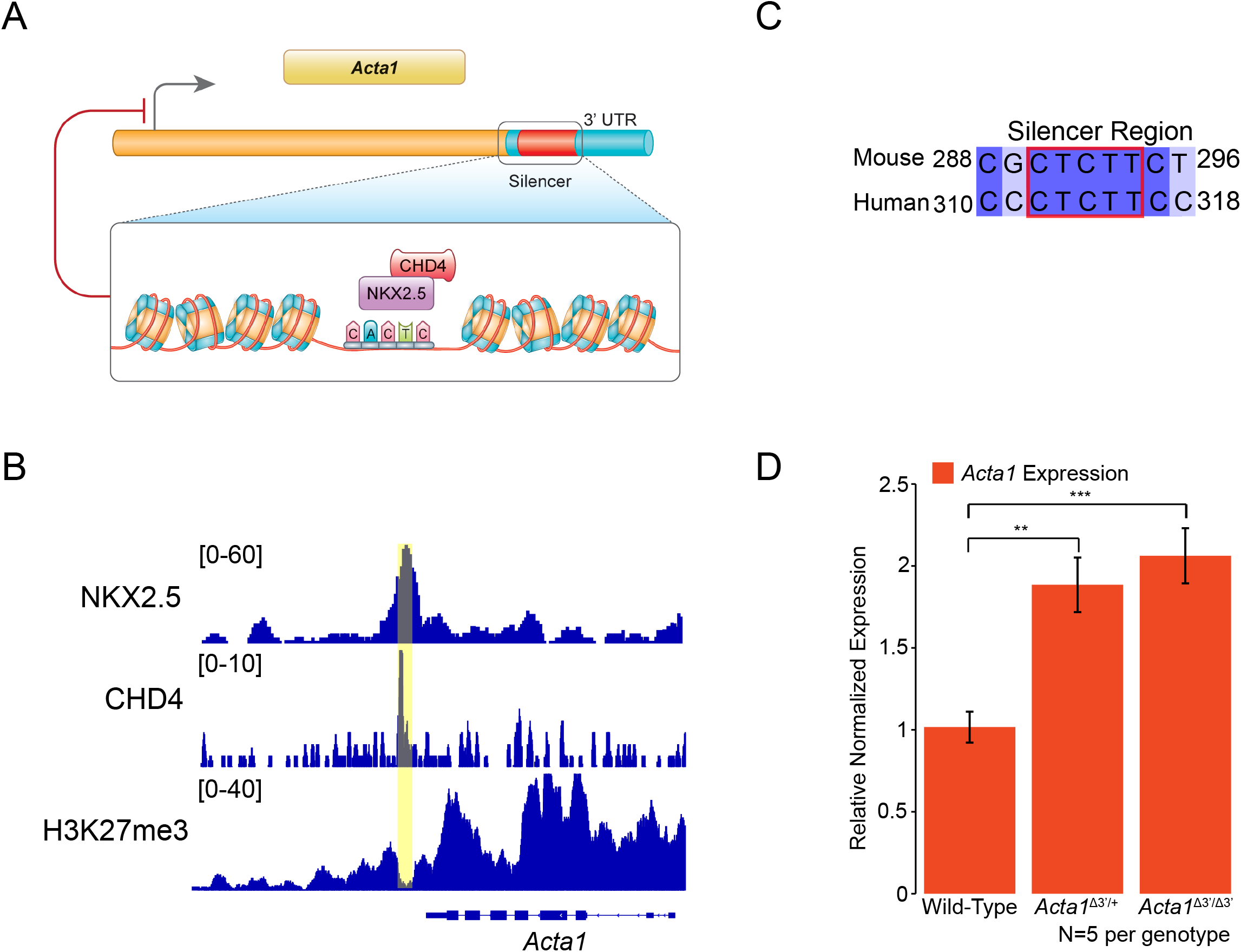
Silencer Downstream of *Acta1* Controlled by CHD4 and NKX2-5 is Required for Repression of *Acta1* at E10.5. (A) NKX2-5-mediated recruitment of CHD4 to target repressor downstream of the TTS in the *Acta1* locus. Putative NKX2-5 motif was identified through motif analysis. (B) Visualization by IGV of ChIP-seq signal across the *Acta1* locus. The repressor region is highlighted with a yellow rectangle. (C) Sequence conservation of CRISPR-deleted region between human and mouse. Red box denotes predicted NKX2-5 binding motif and shows strong conservation with human sequence. (D) Relative expression of *Acta1* in E10.5 embryonic hearts when repressor region excised from one or both alleles. Significance was assessed using an unpaired two-tailed Student’s t test. ** P value < 0.01, *** P value < 0.001, n = 5 non-pooled hearts per genotype. Data normalized to expression of *Pgk1*.

To genetically interrogate whether *Acta1* is repressed in vivo in cardiac tissue thorough this downstream region, we used CRISPR/Cas9 technologies to generate mice in which the putative repressive domain was disrupted (*Acta1^Δ3’^*). These studies reveal that the loss of a single copy of the silent enhancer containing the NKX2-5 binding site in the 3-UTR leads to *Acta1* misexpression in E10.5 hearts in a dominant manner (Figure 4D) with mRNA levels of *Acta1* showing a significant increase relative to wild-type littermate controls (Figure 4D). We conclude that a single copy of an NKX2-5 binding site is sufficient to recruit CHD4 and repress expression of a skeletal actin isoform in the developing mammalian heart. These data further show that, in contrast to its established role in gene activation, NKX2-5 can also mediate cardiac transcriptional repression.

### Deletion of a GATA4-CHD4 site in *Myh11* leads to its mis-expression in developing mouse hearts

We identified a single GATA4 binding site within the third intron of *Myh11*, the gene encoding the smooth muscle myosin heavy chain (SMMHC) protein (Figures 5A, B), an established target of CHD4 (Wilczewski et al. 2018). The canonical GATA4 DNA-binding motif in *Myh11* intron 3 was conserved between mouse and human, co-occupied by CHD4 and GATA4, NKX2-5 and TBX5 during fetal heart development, and flanked by H3K27me3 marked chromatin (Figures 5B, C). Consistent with these observations, we found *Myh11* expression to be significantly increased in GATA4 mutant hearts versus littermate controls (Figure S2). Interestingly, although we detected strong NKX2-5 and TBX5 ChIP signal, the recognition element for NKX2-5 or TBX5 was not detected in either mouse or human sequence. Thus, it may be that GATA4 recruits NKX2-5 and TBX5 as well as CHD4 to this region.

To test the effect of this resion, we used CRISPR/Cas9 technologies to delete the GATA4 binding region in mice (*Myh11^Δi3^*) (Hashimoto et al. 2016). Strikingly, we found that deletion of the *Myh11* silencer caused a large upregulation of *Myh11* in a dominant manner (Figure 5D). *Myh11* expression increased approximately 70-fold in E10.5 heart tissue derived from heterozygous *Myh11^Δi3/+^* mice, whereas in mice homozygous for *Myh11^Δi3/Δi3^*, the deletion increased *Myh11* expression by greater than 200-fold. Collectively these data define a novel silencer of *Myh11* that is regulated by the GATA4-CHD4 complex in association with NKX2-5 and TBX5 and that is essential to repress *Myh11* in the heart.

**Figure 5.**
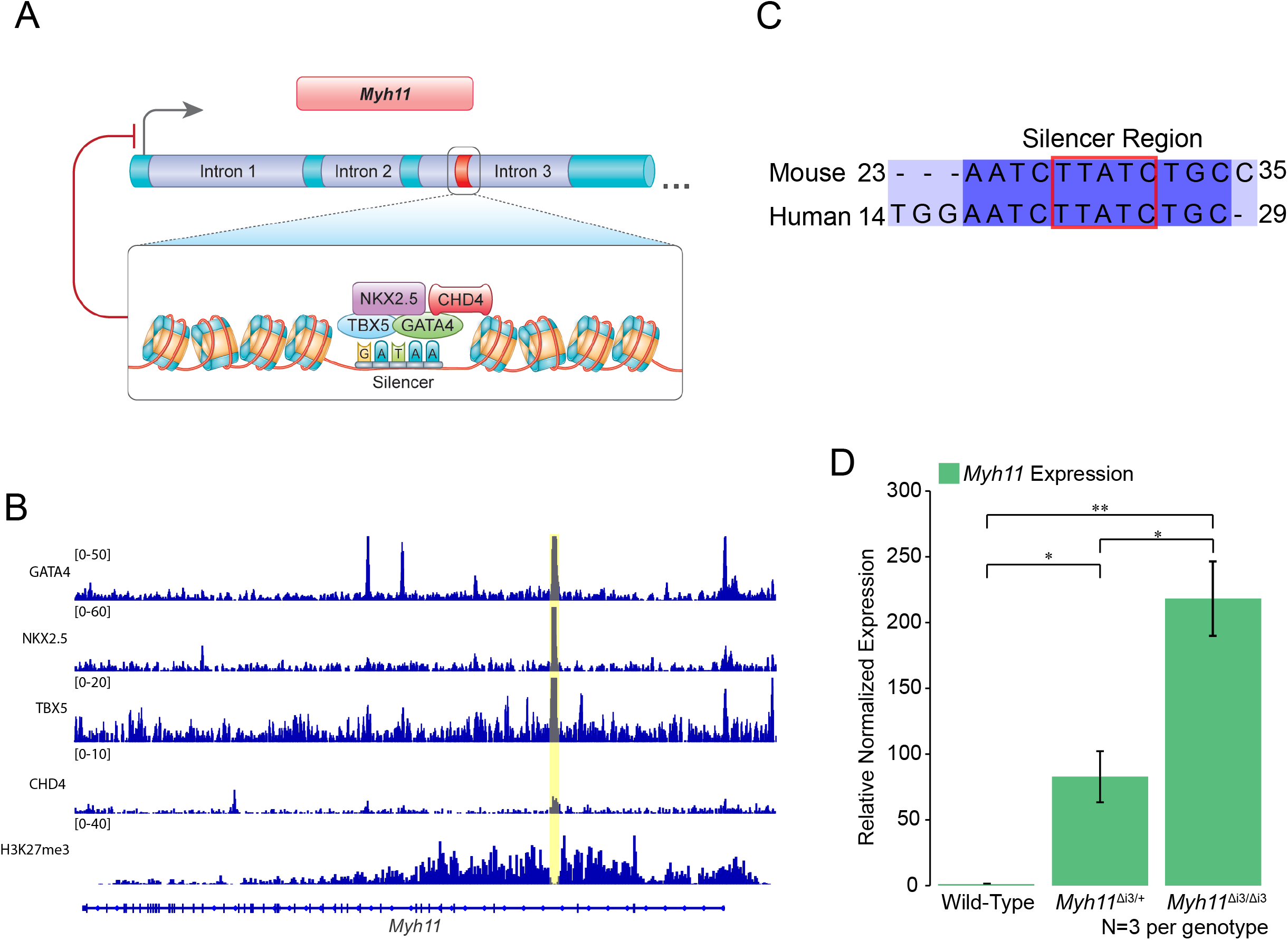
Repressor Region in *Myh11* Controlled by CHD4 and GATA4, NKX2-5, and TBX5 is Required for Repression of *Myh11* at E10.5. (A) Schematic demonstrating GATA4-mediated recruitment of CHD4 to target repressor in the third intron of *Myh11*. A putative GATA4 binding motif was identified through motif analysis using HOMER. (B) IGV Genomic visualization of ChIP-seq signal of of GATA4, NKX2-5, TBX5, CHD4, and H3K27me3 across the *Myh11* locus. The repressor region is highlighted with a yellow bar. (C) Sequence conservation of CRISPR-deleted region between human and mouse. The red box denotes the predicted GATA4 binding motif and shows strong conservation with human sequence. (D) Relative expression of *Myh11* in E10.5 embryonic hearts when the repressor region was excised in one or both alleles. Significance assessed with unpaired twotailed Student’s t test. * P value < 0.05, ** P value < 0.01, n = 3 non-pooled hearts per genotype. Data normalized to expression of *Pgk1*.

## Discussion

Although CHD4 and the NuRD complex are well established to function in repressing transcription, how CHD4/NuRD is recruited to specific loci remained unknown. Here, we demonstrate that the core cardiac transcription factors GATA4, NKX2-5 and TBX5 interact and recruit CHD4 to defined genomic locations associated with H3K27me3. Our findings reveal that NKX2-5, a cardiac transcription factor that has long been understood to function as a transcriptional activator (McCulley and Black 2012), can also function as a cardiac transcriptional repressor. Our studies further show that NKX2-5 and GATA4 function to repress genes incompatible with heart development and function. Because GATA4, NKX2-5 and TBX5 cause a range of congenital heart disease (Basson et al. 1997; Li et al. 1997; Furtado et al. 2017; Steimle and Moskowitz 2017; Zhang et al. 2017; Behiry et al. 2019), our findings imply that the respective patient phenotypes are not only due to loss of cardiac gene expression but also to mis-expression of non-cardiac genes in the developing heart.

Our data supports the hypothesis that mutations in regulatory regions essential for CHD4-mediated repression act in a dominant manner. This may provide one mechanism for the prevalence of CHD in humans with one mutated copy of NKX2-5, Gata4 or Tbx5. Our findings that CHD4 complexes with temporally and spatially regulated cardiac transcription factors further explicates how mutations in the broadly expressed CHD4 protein lead to cardiac specific disease states.

The findings that GATA4, NKX2-5 and TBX5 can recruit CHD4 to the majority of CHD4 repressed target genes in the heart is broadly consistent with studies in B cells demonstrating interaction of CHD4 with lineage-specific transcription factors (Yoshida et al. 2019). However, analyses of GATA4, NKX2-5, TBX5 or the B cell transcription factors fails to uncover any shared sequence homology, conserved sequences or common motif, even at low stringency, thus raising the question of how this diverse set of transcription factors interact with CHD4. Moreover, our analyses of the CHD4 interactome failed to identify any proteins that may function as adapter proteins. Based on these observations, we favor a model by which these sets of transcription factors recognize a secondary structure within CHD4, possibly within or adjacent to the PHD and/or CD domains (Mansfield et al. 2011; Watson et al. 2012; Low et al. 2016; Zhang et al. 2016; Farnung et al. 2020).

CHD4/NuRD is essential for numerous developmental events, such as ensuring proper timing of the switch from stem-cell lineages to differentiated cell types, maintaining cell differentiation, and activating DNA damage response pathways (Larsen et al. 2010; Polo et al. 2010; Scimone et al. 2010; Hosokawa et al. 2013; O’Shaughnessy and Hendrich 2013; Sparmann et al. 2013; Chudnovsky et al. 2014; O’Shaughnessy-Kirwan et al. 2015; Gomez-Del Arco et al. 2016; Zhao et al. 2017; Ostapcuk et al. 2018; Arends et al. 2019; McKenzie et al. 2019; Yoshida et al. 2019; Hou et al. 2020; Sreenivasan et al. 2021). We purpose that one of the generalized functions of CHD4/NuRD is to repress the inappropriate activation of developmental programs of a given tissue or cell type. We suggest that alteration in CHD4 recruitment by tissue specific transcription factors lead to the wide range of CHD4 associated disease states.

## Methods

### Mice

*Chd4^flox/flox^* mice were obtained from Dr. Katia Georgeopolos (Williams et al. 2004). *Nkx2-5^Cre/+^* mice were obtained from Robert Schwartz (Moses et al. 2001). *Chd4* conditional knockout mice and control littermates were obtained by breeding female *Chd4^flox/flox^* mice to male *Chd4^flox/+^; Nkx2-5^Cre/+^* mice. CRISPR/Cas9-mediated genome editing was performed by the UNC Animal Models Core Facility. CRISPR founder mice were bred to wild-type C57Bl6J female mice for two generations. Heterozygous F2 mice were interbred to generate embryos. Research was approved by the Institutional Animal Care and Use Committee at the University of North Carolina and conforms to the Guide for the Care and Use of Laboratory Animals.

### CRISPR/Cas9 Mediated Mouse Engineering and breeding

Cas9 guide RNAs flanking the desired deletion regions were identified using Benchling software. Three guide RNAs at each end of the target sequence were selected for activity testing. Guide RNAs were cloned into a T7 promoter vector followed by *in vitro* transcription and spin column purification. Functional testing was performed by transfecting a mouse embryonic fibroblast cell line with guide RNA and Cas9 protein. The guide RNA target site was amplified from transfected cells and analyzed by ICE (Synthego). One guide RNA at each end of the target sequence was selected and a donor oligonucleotide was included to facilitate homologous recombination to produce a clean deletion event between the guide RNA cut sites. C57BL/6J zygotes were electroporated with 1.2 μM Cas9 protein, 47 ng/ul each guide RNA and 400 ng/ul donor oligonucleotide and implanted in recipient pseudopregnant females. Resulting pups were screened by PCR and sequencing for the presence of the deletion allele. Male founders with the correct deletion were mated to wild-type C57BL/6J females for germline transmission of the deletion allele. The founding *Acta1^Δ3’^* and *Myh11^Δi3^* males were bred to wildtype mice for two generations, and the genotypes of *Acta1^Δ3’^* and *Myh11^Δi3^* founding males and all F2 offspring were confirmed by sequencing and PCR/restriction digests. F2 mice were intercrossed and hearts derived from homozygous, heterozygous, and wildtype littermates were collected and assayed for *Acta1* and *Myh11* mis-expression, respectively.

### Motif Discovery

De novo motif discovery on CHD4 chromatin immunoprecipitation followed by high-throughput sequencing (ChIP-seq) regions (Wilczewski et al. 2018) (GSE109012) from embryonic day (E)10.0 cardiac tissue was performed using Hypergeometric Optimization of Motif EnRichment (HOMER) (Heinz et al. 2010).

### CHD4 immunoaffinity purification

E10.5 wild-type CD1 hearts (minimum twenty-eight hearts per IP, three separate biological replicates) were harvested in cold PBS, snap frozen and cryolysed as previously described (Kaltenbrun et al. 2013). Frozen tissue powder was resuspended in optimized lysis buffer (5ml/1g powder) (20mM K-HEPES pH7.4, 0.11M KOAC, 2mM MgCl2, 0.1% Tween 20, 1μM ZnCl2, 1mM CaCl2, 0.5% Triton-X 100, 150mM NaCl2, protease inhibitor (Sigma), phosphatase inhibitor (Sigma)). CHD4 complexes were solubilized and immunoaffinity purified as previously described (Cristea and Chait 2011; Kaltenbrun et al. 2013) using rabbit anti-CHD4 antibody (Abcam #ab72418) or negative control custom rabbit anti-GFP antibody (Cristea et al. 2005) with elution at 95°C for 10 minutes. The immunoisolated proteins were resolved (~ 1 cm) by SDS-PAGE, and visualized by Coomassie blue. Each lane was subjected to in-gel digestion with trypsin as previously reported (Waldron et al. 2016).

### (Parallel Reaction Monitoring) PRM nLC-MS/MS

Immunoisolated proteins were resolved (~ 1 cm) by SDS-PAGE, and visualized by Coomassie blue. Each lane was subjected to in-gel digestion with trypsin as previously reported (Waldron et al. 2016). Desalted peptides (2μl each) were analyzed by nano-liquid chromatography (nLC)-MS/MS using a Dionex Ultimate 3000 nRSLC system directly coupled to a Q-exactive HF orbitrap mass spectrometer (ThermoFisher Scientific) equipped with an EASY-Spray ion source (Thermo, Fisher Scientific). Peptide mixtures were evaporated in vacuo and resuspended in 1% Trifluoroacetic acid/4%acetonitrile/95% water and loaded onto an 50 cm long column with a 75 um internal diameter (PepMap) and separated over a 60 min gradient with a mobile phase containing aqueous acetonitrile and 0.1% formic acid programmed from 1% to 3% acetonitrile over 12 min, then 3 to 35% acetonitrile over 60 min, then 35 to 97% acetonitrile over 1 min, followed by 10 min at 97% acetonitrile and 97 to 70% over 1 min, all at a flow rate of 250 nL/min.

The PRM analysis was carried out as previously reported(Justice et al. 2021). Briefly, the method consisted of targeted MS2 scans at a resolution of 60,000 and with an AGC value of 1×10^6^, a max injection time of 500 ms, a 0.8 *m/z* isolation window, a fixed first mass of 150 *m/z*, and normalized collision energy of 27, which were recorded as profile data. The targeted MS2 methods were controlled with a timed inclusion list containing the target precursor *m/z* value, charge, and a retention time window that were determined from shotgun analysis.

### cDNA synthesis and qPCR

RNA from individual E10.5 hearts (N=3 or 5 per genotype) was isolated using the RNAqueous-Micro Total RNA Isolation Kit following manufacturer recommended protocols (Invitrogen). cDNA synthesis was performed using random hexamers and SuperScript IV reverse transcriptase (Invitrogen). Quantitative PCR was performed using PowerUP SYBR Green Master Mix (ThermoFisher) at standard cycling conditions on a QuantStudio 6 Flex instrument (ThermoFisher) with the following primers:

*Myh11* F1: GCTAATCCACCCCCGGAGTA
*Myh11* R1: TCGCTGAGCTGCCCTTTCT
*Acta1* F1: GTGACCACAGCTGAACGTG
*Acta1* R1: CCAGGGAGGAGGAAGAGG
*Pgk1* F1: GTCGTGATGAGGGTGGACTT
*Pgk1* R1: AAGGACAACGGACTTGGCTC

### Proximity ligation assay (PLA)

A standardized procedure for PLA was used (Jalili et al. 2018). Briefly, HEK-293 cells were seeded on circular coverslips and transfected with specific plasmids for 72 hours using a 3:1 ratio of PEI to pDNA. Cells were fixed with 4% paraformaldehyde for 10 minutes, followed by permeabilization and blocking (10% heat-inactivated goat-serum, 1% Triton-X 100) for 1 hour. Cells were then probed with two specific primary antibodies raised in different species, mouse anti-TurboGFP (1:250, Origene, TA150041) / rabbit anti-Flag (1:500, Sigma, F7425) overnight at 4 °C. The cells were incubated with PLA probes for 1 hr at 37 °C, with ligase for 30 min at 37 °C and with polymerase for 100 min at 37 °C, based on manufacturer protocols. Cells were stained with DAPI and then mounted with PermaFluor mounting medium (ThermoScientific TA030FM). Images were captured on a Zeiss LSM 700 laser scanning confocal microscope.

### DNA Constructs

Full length human *CHD4* tagged at C-terminus with Flag/Myc construct was obtained from Origene (RC224232). Full-length human *GATA4* cDNA was amplified with 5’ primer (ATTAGCGATCGCCATGTATCAG) and 3’ primer (CGTACGCGTCGCAGTGAT). Amplicons were digested with restriction enzymes AsiSI / MluI and inserted into pCMV6-AC-turboGFP vector (Origene, PS100010). Full-length human *NKX2-5* cDNA was amplified with 5’ primer (ATTAGGATCCATGTTCCCCAGCCCTG) and 3’ primer (GTCGACTCACCAGGCTCGGATACCAT). Amplicons were digested with restriction enzymes AsiSI / MluI and inserted into pCMV6-AC-turboGFP vector (Origene, PS100010).

### NGS Datasets Used

GATA4 Genome Occupancy at E12.5 in the developing mouse heart (GEO:GSE52123) (He et al. 2014).

NKX2-5 Genome Occupancy at E12.5 in the developing mouse heart (GEO:GSE124008) (Akerberg et al. 2019)

TBX5 Genome Occupancy at E12.5 in the developing mouse heart (GEO:GSE124008) (Akerberg et al. 2019)

CHD4 Genome Occupancy at E10.0 in the developing mouse heart (GEO:GSE109012) (Wilczewski et al. 2018)

CHD4 Transcriptomics at E10.5 in the developing mouse heart (GEO:GSE109012) (Wilczewski et al. 2018)

H3K27me3 Histone Modification at E10.5 in the developing mouse heart (GEO:GSE86693) (Consortium 2012)

### Genome Annotation and Co-Occupancy Analysis

BAM and BED files were obtained from the Gene Expression Omnibus aligned to mm10. Peaks were called using MACS2 (v2.2.7.1) (Zhang et al. 2008) using default settings with a q-value of 0.01. High confidence peaks appearing in both replicates were retained for downstream analysis. Peak annotation was conducted with HOMER (v4.10) (Heinz et al. 2010). Peaks were annotated using the ChIP-Seeker (v1.24.0) R package(Yu et al. 2015). Biological Process GO terms for pe ak regions were generated using R package ClusterProfiler (Qian et al. 2012). Gene tracks were visualized using IGV (Robinson et al. 2011). Peak overlaps were determined using BedTools, an d the upset plot was generated using UpSetR v1.4.0(Conway et al. 2017) in R. Overlaps between CHD4, GATA4, NKX2-5, TBX5, and H3K4me3 were plotted in relation to their chromosomal location using the karyoploteR (v1.14.0) R package(Gel and Serra 2017).

## Competing interest statement

The authors declare no competing interests.

## Acknowledgements

This work was supported by grants R01HL156424 NIH/NHLBI to F.L.C, R01HD089275 NIH/NHLBI to F.L.C. and I.M.C., and 2UM1HL098166 to W.T.P.

## Author contributions

Z.R., W.S., and F.C. designed most of the experiments. Z.R., W.S., and C.W. performed RNA-seq, B.A. performed the ChIP-seq. Z.R., W.S., L.W., C.W., A.H., and B.A. performed and interpreted the bioinformatic analyses. Z.R., C.W, and X.S. performed the IP/PRM-MS experiments and analyses. Z.R. and W.S. generated the knockout mouse lines and performed the RT-qPCR. W.P., I.C., I.D., and F.C. supervised the project. Z.R., W.S., L.W., and F.C. wrote the manuscript with feedback from all authors.

---

**Figure S1.**
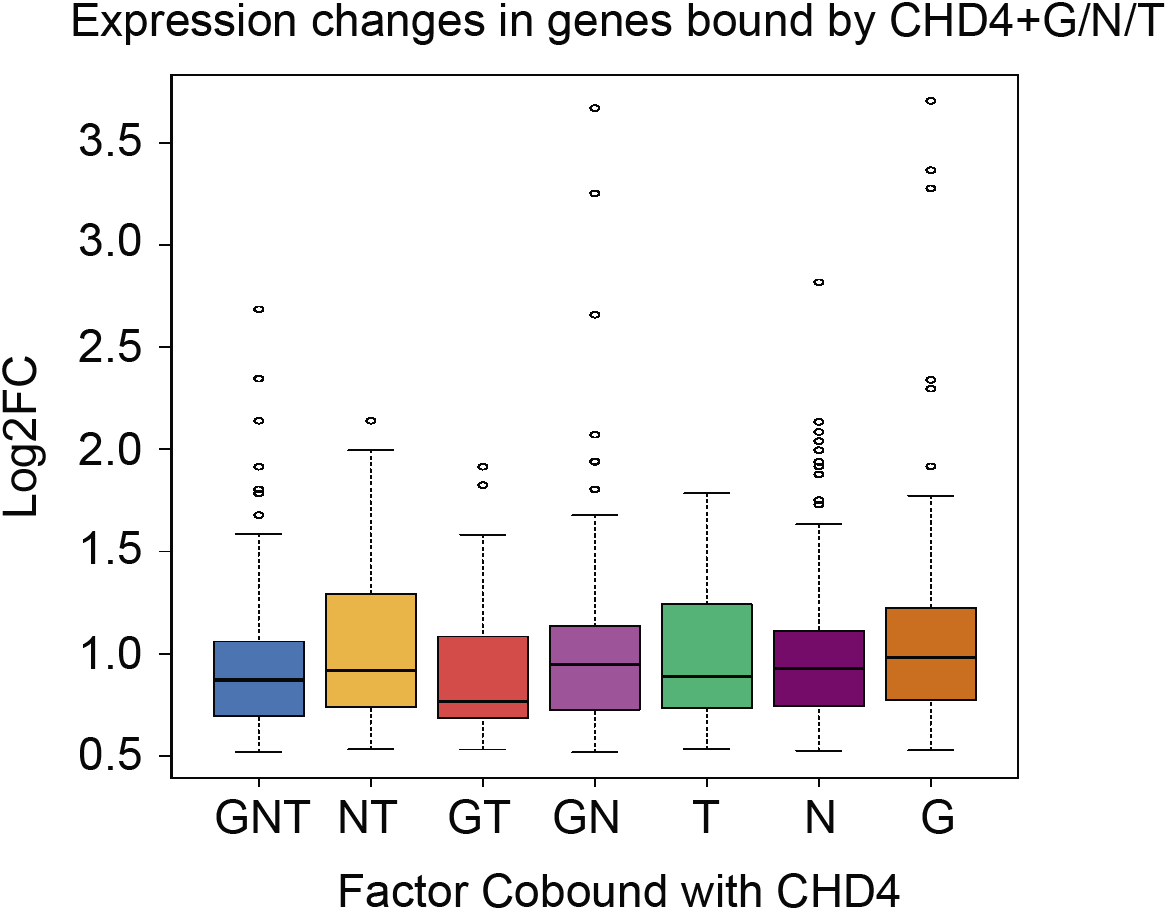
Change in expression of significanly repressed genes at E10.5 in CHD4 conditional knockout hearts. Genes were grouped by CHD4 association with one or a combination of GATA4, NKX2-5 and TBX5. Data is presented as median log2 (Fold-Change) and interquartile ranges.

**Figure S2.**
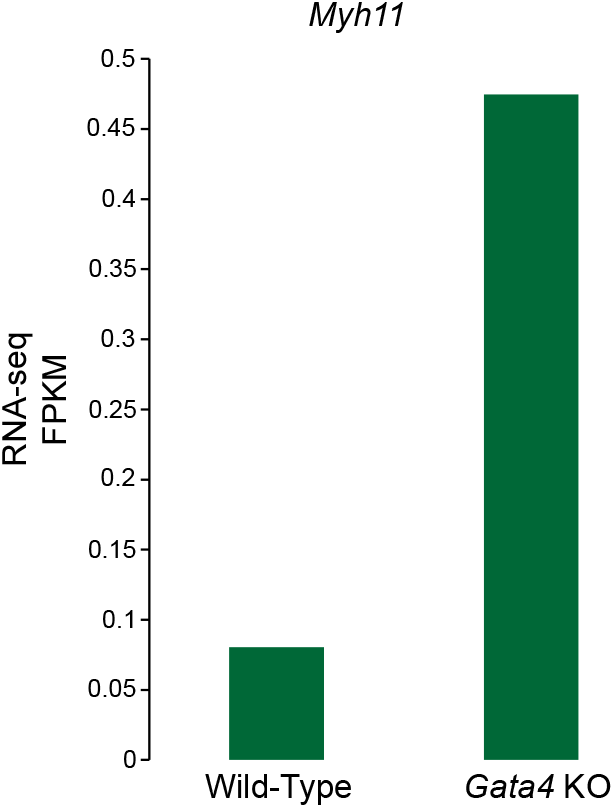
Fragments Per Kilobase of transcript per Million (FPKM) mapped reads of *Myh11* in RNA-seq of *Gata4 KO* hearts.

## Notes

### Competing Interest Statement

The authors have declared no competing interest.

